# Distinct Long-term Effects of Precision X-Radiation on Reflex Saliva Flow Rate and Tissue Integrity in a Preclinical Model of Chronic Hyposalivation

**DOI:** 10.1101/2020.05.26.117937

**Authors:** Syed Mohammed Musheer Aalam, Ishaq A. Viringipurampeer, Matthew C. Walb, Erik J. Tryggestad, Chitra P. Emperumal, Jianning Song, Xuewen Xu, Rajan Saini, Isabelle M.A. Lombaert, Jann N. Sarkaria, Joaquin Garcia, Jeffrey R. Janus, Nagarajan Kannan

## Abstract

Chronic salivary hypofunction and xerostomia are common side effects of radiation therapy which is an essential component in the curative management in patients with head & neck cancers. Over the years, improvements in delivery techniques such as image-guided intensity modulated radiation therapy have led to improvement in cancer management but chronic hyposalivation continues to be a challenge that causes long-term health implications resulting in compromised quality of life. Recent advances in salivary stem cell research promise new frontier in the treatment of radiation-induced hyposalivation by initiating regeneration of radiation-damaged salivary parenchymal cells. Lack of a standard preclinical immunodeficient model to assess radiation-induced changes objectively and quantitatively in salivary flow rates will impede rapid progress towards the development of cellular therapies for chronic salivary dysfunction and attendant xerostomia. Herein, we report the first fully characterized novel cone-beam computed tomography (CBCT)-guided precision ionizing radiation (IR) induced chronic hyposalivation model in radiosensitive, immunodeficient transgenic NSG-SGM3 mice expressing three human cytokines including c-KIT ligand/stem cell factor. Additionally, we also report a novel and instantaneous method to objectively assess the kinetics of pilocarpine-stimulated salivary flowrate. Comprehensive structural and functional characterization of salivary glands revealed previously unknown and highly complex gender, age, IR dose and salivary gland subtype-specific effects of salivary-ablative precision IR.

## Introduction

Head and neck cancer accounts for ~4% of all cancers in the United States (1). The current standard of care for management can involve surgery, radiation therapy, chemotherapy, targeted therapy or a combination of these treatment modalities depending on various factors including the stages of cancer, the surgical accessibility of the tumor, and morbidity associated with each modality (2). In nearly 70% of the patients undergoing radiation therapy due to location of many of these oral cancers, non-diseased adjacent tissues such as salivary parenchyma is irreparably damaged due to its high sensitivity to radiation. This often results in radiation-induced ‘objective’ hyposalivation or a ‘subjective’ clinical condition referred to as xerostomia or dry mouth (2, 3). Complications of long-term radiation induced hyposalivation include impaired taste, difficulties in speech, mastication and denture retention, increased risk of dental caries and an overall compromised quality-of-life (QOL) (4). Management of hyposalivation often involves the use of cholinergic drugs such as pilocarpine (5) or cevimeline (6). The effectiveness of these systemic sialagogues often depends on the presence of functionally-intact glandular tissue units (5). Recently, putative stem cells, that can restore the salivary gland functions upon orthotopic transplantation in radioablated salivary glands of recipient mice, have been reported in the submandibular glands of mice (7–9) and humans (10). To compare the quality and quantity of salivary regenerative units in transplant experiments, a standardized *in vivo* assay must be developed. However, there are several limitations with currently available models of hyposalivation. 1) Their derivation involves IR and there is no consensus on sources of IR, radiation fields, doses or treatment protocols. 2) A reliable method to measure kinetics of stimulated reflex saliva is lacking. 3) A comprehensive characterization of cellular components of radioablated salivary glands in an immunodeficient mouse model of chronic hyposalivation is lacking. 4) Human salivary stem cells are suggested to express c-KIT receptor, which interacts poorly with murine c-KIT ligand and therefore mouse models lacking the expression of human cKIT ligand may not fully support the engraftment of human salivary stem cells (10). These lacunae need to be addressed before such preclinical models could be routinely used for human salivary stem cell evaluation. Therefore, in the present study, based on comprehensively characterization of precision irradiated male and female (young and old) immunodeficient transgenic NSG-SGM3 mice expressing human c-KIT ligand, we have identified suitable models for chronic hyposalivation studies for preclinical assessment of experimental therapies. Further, we demonstrate Schirmer’s test as an accurate, reliable method to assess quantitative changes in saliva flow rate in real time in our hyposalivation model. Moreover, characterization of salivary glands at necropsy further revealed highly complex gender, age, IR dose and salivary gland subtype-specific effects of salivary-ablative precision IR in NSG-SGM3 mice. In summary, we have developed the first humanized mouse model that mimics a challenging, chronic lifestyle problem in cancer survivors, which provides a robust platform to enable studies to measure regenerative outcome following patient-derived salivary cell transplants.

## Results and Discussion

To generate a preclinical model of chronic hyposalivation here in, we utilized 3D image guided stereotactic X-RAD SmART+small animal irradiator system (Precision X-ray Inc., North Bradford, CT) to radioablate the major salivary glands of immunodeficient NSG-SGM3 (NOD.Cg-*Prkdc^scid^ Il2rg^tm1Wjl^* Tg (CMV-IL3, CSF2, KITLG) mice (11). Details of the functionality of this irradiator are described elsewhere (12–14). Dosimetric calculations were performed with an in-house 1D “point dose calculator” (PDC) tool developed by our medical physics group to enable efficient dose calculations (single prescription reference point on the central beam axis) (15). This simple 1D PDC was verified using the vendor supplied 3D Monte Carlo treatment planning system, SmART-ATP version 1.1 (SmART Scientific Solutions B.V., Maastricht, Netherlands), which is based on the open-source Monte Carlo code (EGSnrc/BEAMnrc)(16, 17) (Fig1A&B). The chosen beam arrangement was parallel opposed beams using a 10 mm circular collimator, with tipped laterals (89° and 271°) aligned to match beam divergence along the brain edge in order to maximize brain sparing and achieve specific targeting of major salivary glands along the midline of the mouse in the neck region (Fig1C (i-iii)). The individual young (~21 weeks) male or female or old (~53 weeks) female mouse was restrained on a bite block using an anesthesia nose cone and imaged with cone beam computed tomography (CBCT). Subsequently, CBCT image-guidance defined target and single radiation dose ranging from 0-7.5 Gy for young and old mice was delivered at target site i.e., major salivary glands. Both sham and precision IR treated mice were followed for 6 months. The young mice precision irradiated with 5 Gy showed transient alopecia whereas, higher dose of 7.5 Gy showed permanent alopecia in the mandibular region (Fig1D). However, we did not observe any significant change in body weight of young or old mice (Fig1E) throughout the study period or precision IR related mortality (Fig1F), which suggests that both male and female *Prkdc^scid^* mutant highly radiosensitive NSG-SGM3 mice are able to tolerate Sub-lethal dose is 3.5 Gy (18) of IR when delivered in a stereotactic manner on the major glands.

**Figure 1.**
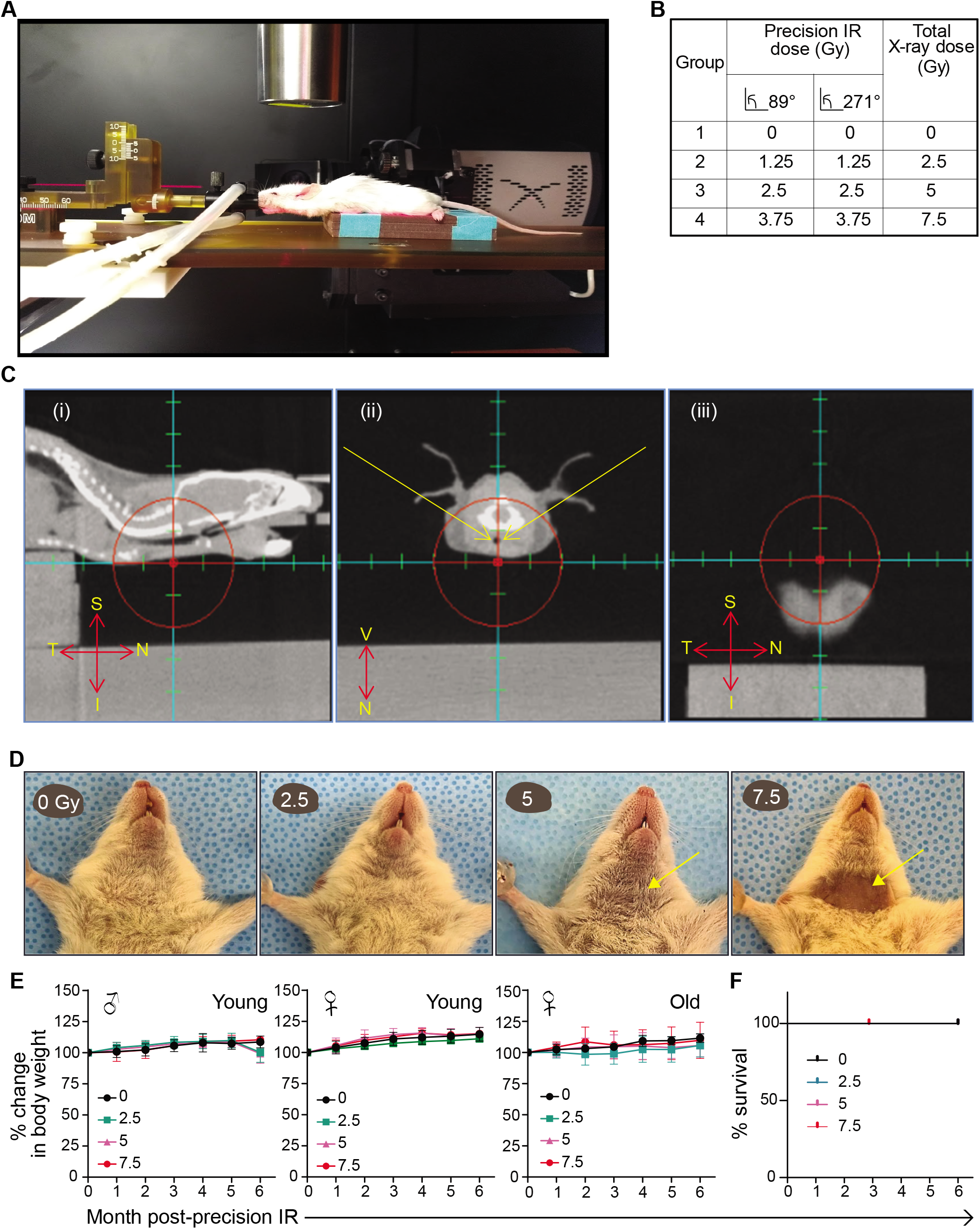
Effects of CBCT-guided stereotactic salivary-ablative precision IR on aging highly immunodeficient and partially humanized NSG-SGM3 mice. A) A photograph showing a dummy mouse restrained on stereotactic stage of the X-RAD SmART irradiator. B) Dosimetry and angle of delivery calculated using vendor supplied Monte Carlo treatment planning system. C) Representative images showing CBCT-guided target placement at the midline of the mouse neck region; (i) axial, (ii) sagittal, and (iii) temporal views. D) Representative head and neck image highlighting chronic alopecia in neck region associated with 0, 2.5, 5 and 7.5 Gy IR (female; n=5 for each group). E) Body weight (in grams) and age (in months) of NSG-SGM3 mice used in this study; male (n=19), female (n=20) and old female (n=20). F) Long-term survival plot in young mice showing lack of precision radiation associated mortality (male n=19; female n=20).

Salivary stimulation and secretion is a nerve-mediated reflex action modulated by the central nervous system in response to a stimulus, and this activity is an important indicator of functional health of salivary glands (19). In hyposalivation animal models, cholinergic drugs such as pilocarpine or carbachol are often used to stimulate saliva secretion (10, 20, 21). Although, there are several methods that have been reported to collect stimulated reflex saliva and evaluate changes in salivary flow rate in vivo or ex vivo (Supplementary table1), the field lacks an operator independent and objective method for instantaneous kinetic measurement of stimulated saliva flowrate in vivo. Therefore, we adopted Schirmer’s test strip method to assess quantitative changes in salivary flow rate in NSG-SGM3 mice. Schirmer’s tests are used in clinical practice to measure eye dryness (22). We tested this method against the most commonly used gravimetric method. We injected 2 mg/kg of pilocarpine sub-cutaneously (SubQ) into the dorsal flanks of mice and started continuously collecting saliva 5 minutes post-injection from the floor of mouth using pre-weighed filter paper or Schirmer’s test strips for 30 minutes. The filter paper or Schirmer’s test strips were changed every five minutes and saliva migration under the capillary action (mm) was recorded. Subsequently, weight of blotted strips was determined gravimetrically (Fig2A). Schirmer’s test demonstrated gender-specific and age-specific differences in stimulated reflex saliva flow rate captured over 30 minutes. The saliva secretion peaks at 10 minutes after stimulation in all mice irrespective of gender, age or radiation treatment and declines thereafter (Fig2B). Sexual dimorphism was reported in submandibular gland (SMG) in mouse models with respect to the unique presence of granular convoluted tubular structures in male mice (23). We found such structures in SMG of male NSG-SGM3 only (Supplementary Fig 1). Additionally, Schirmer’s test distinguished young female mice having significantly lower initial stimulated saliva flowrates compared to young male and old female mice at −1 weeks and +3 weeks of sham-irradiation (Fig2B). Notably, the old female mice exhibited relatively more rapid secretory loss after 10 min (Fig2B). This trend was consistent during the 6-7 month measurement period in sham irradiated mice. Furthermore, various measurements of saliva fraction over the period of 24 weeks in young male and female mice demonstrated that our Schirmer’s test strip method was highly correlated with gravimetric method in both male (R2=0.9031, p<0.0001) and female (R^2^=0.7289, p<0.0001) NSG-SGM3 mice (Fig2D). Taken together, our results suggest that Schirmer’s test strips offer a, rapid and reliable quantitative method to assess real-time change in saliva flow rate in mice. This method can be easily adapted to study saliva flow rate in other mouse strains and species. Moreover, Schirmer’s test strips circumvent the variability associated with salivary collection and measurement methods. In fact, in our study the saliva measurement with Schirmer’s test strips was found to be consistent between three independent operators (data not shown). Salivary gland hypofunction resulting in chronic hyposalivation is often one of the consequences associated with radiotherapy of head and neck malignancies. Patients with such complications experience QOL or oral health issues (4). Modeling chronic radiation-induced hyposalivation in animals has been challenging, as there is no consensus on radiation dose, gender, age and duration of the study period. Therefore, we systematically characterized stereotactic radiation dose-dependent chronic reflex hyposalivation in preclinical NSG-SGM3 model. We determined the kinetics of pilocarpine stimulated reflex saliva flow rates over the course of 30 minutes using Schirmer’s test strips in young male, female and old female mice irradiated with 0, 2.5, 5 and 7.5 Gy doses. At 24 weeks post-irradiation, we observed significant delay in the onset of stimulated reflex saliva secretion and reduction in salivary flowrates in mice irradiated with radiation doses of 5 and 7.5 Gy compared to 0 Gy controls (Fig3A). However, mice irradiated with 2.5 Gy dose showed a marginal decline in stimulated reflex saliva flowrate compared to 0 Gy controls in young male and female mice (Fig3A). Moreover, at early time points (5 and 10 min) following pilocarpine stimulation, salivary flow rate is significantly different between young male and female mice at the high doses (5 and/or 7.5 Gy) (pvalue < 0.0055 and 0.0023) (Fig3B). Further, to assess chronic hyposalivation in our pre-clinical NSG-SGM3 model, we continued the measurement of stimulated reflex saliva flow rate until a period of 6 months post-irradiation. We observed marked reduction in stimulated reflex saliva fraction immediately following irradiation in young male, young female and old female mice. After 2 months post-irradiation with 5 and 7.5 Gy doses, the stimulated reflex saliva flow rate stabilized at 50% and was sustained until the endpoint of 6 months relative to 0 Gy controls (Fig3C&D). Interestingly, mice irradiated with 2.5 Gy dose showed initial decline in stimulated reflex saliva flow rate, but this was followed by restoration as early as 3 weeks after irradiation although not to the extent similar to that of 0 Gy controls (Fig3C&D). This restoration of salivary flowrate could be partially attributed to ongoing salivary stem cell activity that may contribute to functional recovery of the gland. Interestingly, starting at 2 months (+6 week and +24 week post-IR), the fraction of saliva in young females was significantly lower compared to young male mice (Fig3D).

**Figure 2.**
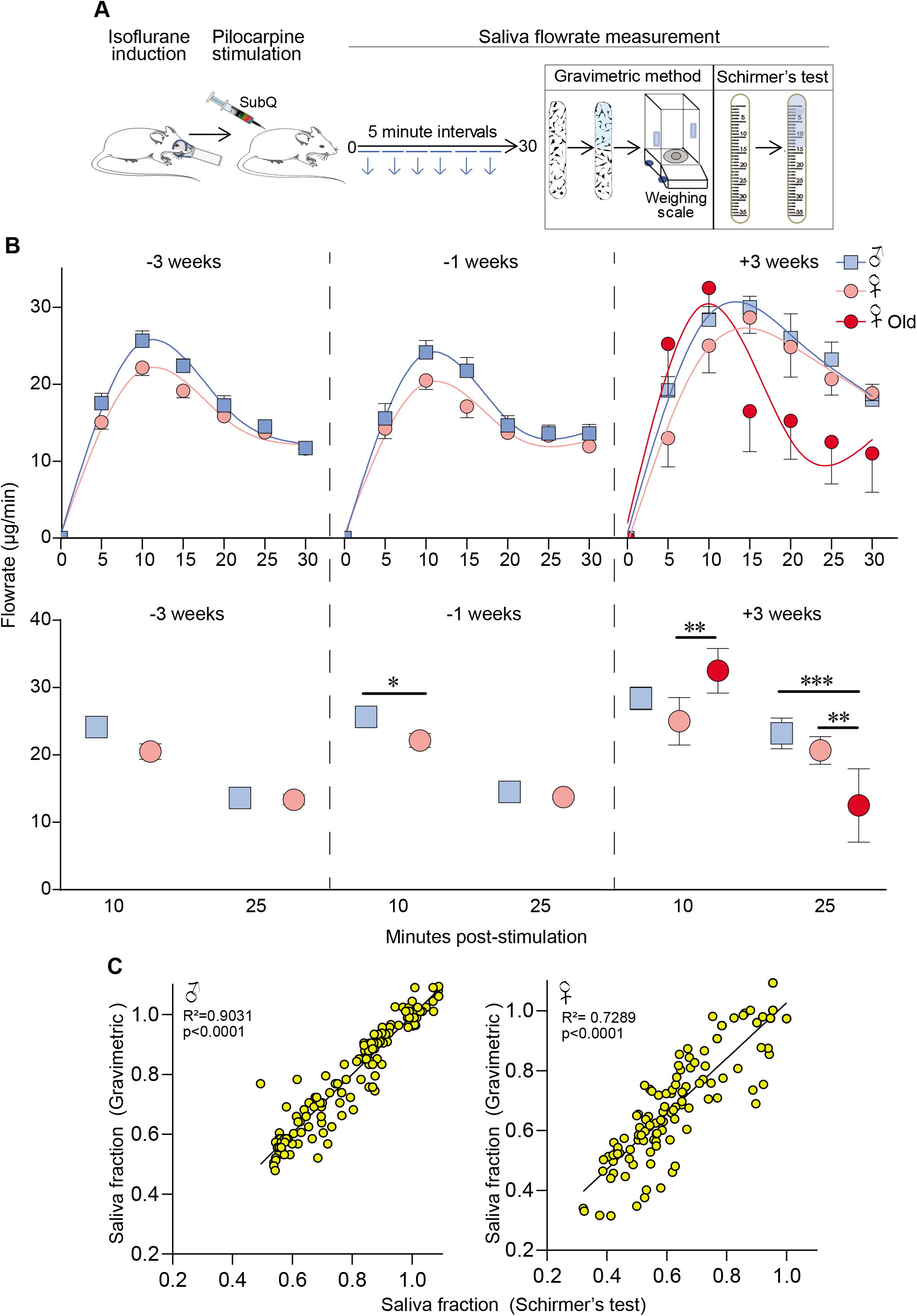
Objective and instantaneous assessment of pilocarpine-stimulated reflex saliva flow in preclinical NSG-SGM3 model. **A**) Illustration of workflow to measure pilocarpine-stimulated reflex saliva flow by Schirmer’s test strips and gravimetry. **B**) Kinetics of pilocarpine-stimulated reflex saliva flowrate (μg/min) in NSG-GSM3 mice (male (n=5), female (n=5) and old female (n=4). **C**) Barplot showing gender and age differences in saliva flowrate at 10 and 25 mins post-stimulation male (n=5), female (n=5) and old female (n=4). **C**) X-Y plot showing correlation between gravimetric vs Schirmer’s test method based detection of pilocarpine-stimulated reflex saliva fraction in male and female mice (male (n=19); female (n=20); *denotes p<0.01.

**Figure 3.**
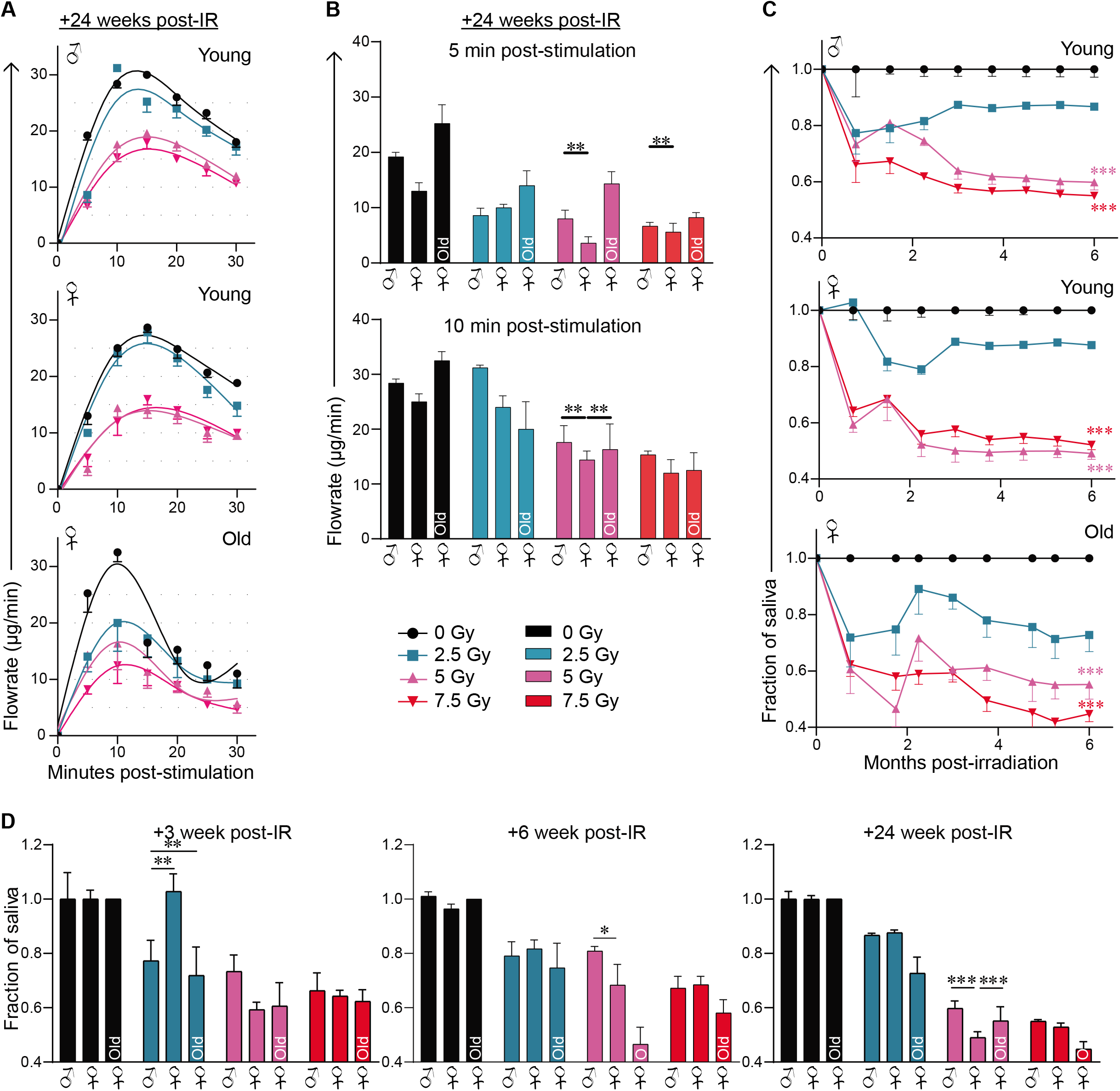
Effects of precision IR doses on chronic hyposalivation in preclinical NSG-SGM3 model. **A**) Kinetics of pilocarpine-stimulated reflex saliva flow rate (μg/min) in mice treated with different precision IR doses (male (n=5 all groups except for 7.5Gy IR dose n=4); female n=5 or n=4 old female for each experimental group). **B**) Gender -and age-specific significant differences in pilocarpine-stimulated reflex saliva fraction in mice treated with different precision IR doses. Data is relative to respective sham treated controls (male (n=5 all groups except for 7.5Gy IR dose n=4); female n=5 or old female n=4 for each experimental group). **C**) High-dose precision irradiated mice displaying delayed onset of stimulated reflex saliva flowrate in time (0, 2, 4, 6 months) (male (n=5 all groups except for 7.5Gy IR dose n=4; female n=5 or old female n=4 for each experimental group). **D**) Barplot showing significant changes in saliva fraction at +3, +6, and +24 weeks in irradiated young male, female and old female mice relative 0 Gy controls (male (n=5 all groups except for 7.5Gy IR dose n=4; female n=5 or old female n=4 for each experimental group.*p<0.01; **p<0.001 and ***p<0.0001

Radiation-induced salivary gland injury is also associated with glandular shrinkage, loss of acinar cell area, inflammation, fibrosis, microvascular injury and atrophy (24). Therefore, at the endpoint of 24 weeks, we sacrificed sham and irradiated mice and performed neck dissection to harvest salivary gland tissue. Overall the sham-irradiated whole salivary glands were significantly larger in old female mice than young female mice relative to their body weight (Fig4A). At necropsy, we observed glandular shrinkage irrespective of gender and age in cohorts of mice irradiated with 5 and 7.5 Gy doses albeit glandular shrinkage was not evident with 2.5 Gy dose in comparison to 0 Gy controls (Fig4B-C). Consistently, we observed significant dose dependent reduction in salivary gland weight in young and old female mice (Fig4C). Interestingly, we did not observe any correlation between radiation doses and salivary gland weight in young male mice due to variability across the radiation doses (Fig4C). Further, we prepared tissue sections from harvested parotid, submandibular and sublingual glands of young female mice irradiated with 0 and 7.5 Gy doses and performed histological and immunohistochemical analysis. Hematoxlin and Eosin (H&E) staining revealed gross changes in tissue morphology and loss of acinar and ductal cells in all major salivary glands of irradiated mice when compared to controls (Fig4D-i). Similarly, Sirius red (Fig4D-ii) and trichrome (Fig4D-iii) staining showed thickening of collagen fibers and fibrosis in tissue sections obtained from irradiated mice when compared to controls. Moreover, to assess the extent of radiation induced damage to salivary epithelium, we stained whole salivary tissue sections for epithelial cell marker EpCAM by immunostaining (25). We observed reduced expression of EpCAM in salivary epithelium of irradiated mice compared to controls, which could be attributed to the loss of functional ducts and acini (Fig4D-iv). To validate this observation we performed immunostaining for Aquaporin 5 (Aqp5), a membrane protein that facilitates water movement across the basal/lateral/ apical membrane in acinar cells (26). Consistently, our results revealed reduction in expression of Aqp5 in acinar cells of irradiated mice compared to controls (Fig4D-v). Interestingly, we observed higher expression of β-catenin in ductal cells of irradiated mice relative to controls (Fig4D-v), suggesting ongoing tissue remodeling activity (25, 27, 28). In addition, we performed immunostaining of NKCC1, a cotransporter highly expressed in baso-lateral membrane of acinar cells. We observed reduced expression of NKCC1 in submandibular glands of irradiated mice when compared to controls (Fig4D-vi). However, we did not observe any changes in NKCC1 expression in parotid and sublingual glands in irradiated and control mice.

**Figure 4.**
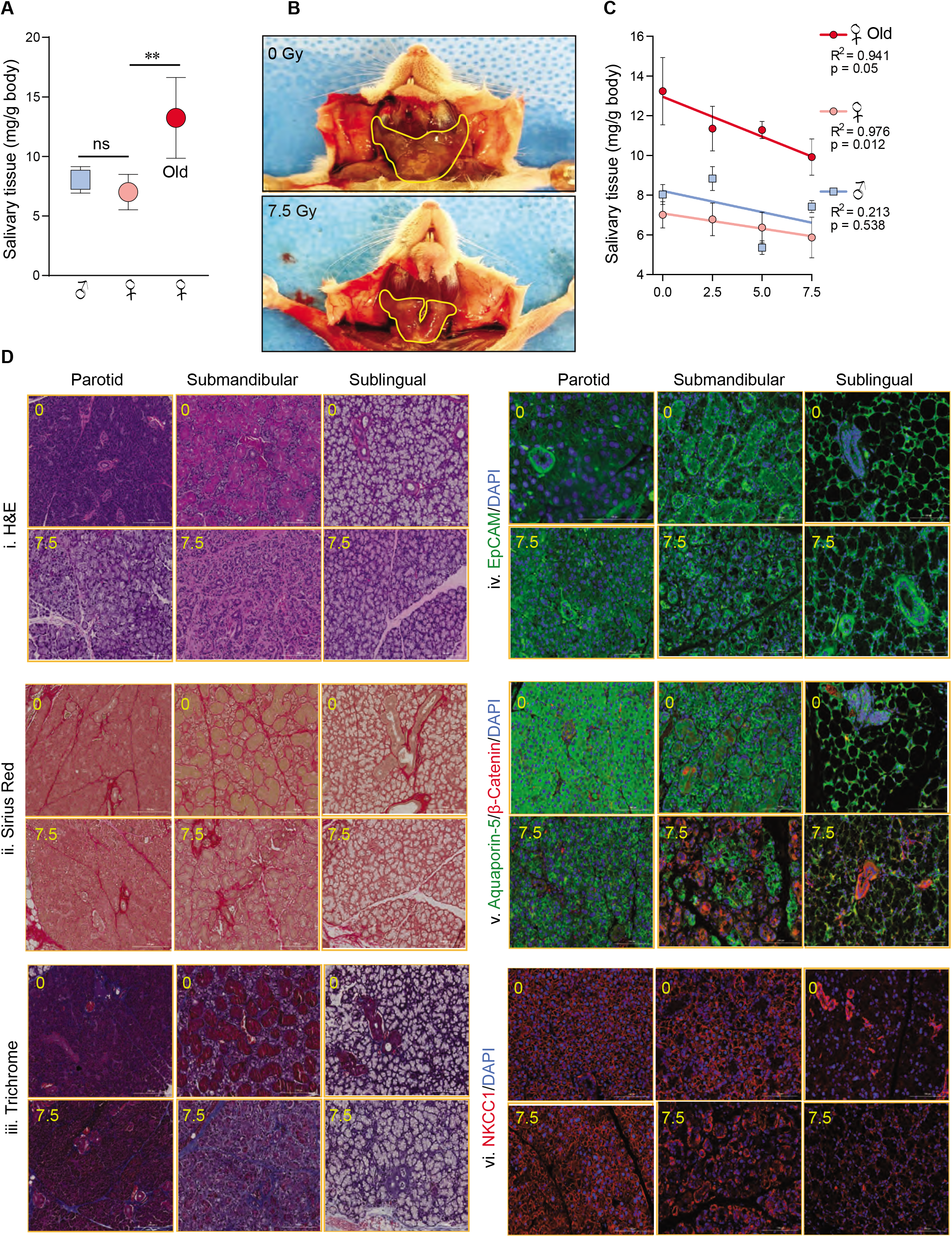
Assessment at necropsy reveals 6 month post-precision IR associated changes on salivary tissue, stroma, epithelial cell polarity, and integrity in preclinical NSG-SGM3 model. **A**) Barplot showing baseline differences in wet weight of total salivary gland tissue (mg/g body weight) obtained from mice at the time of necropsy (male n=19; female n=20 or old female n=20. **B**) Representative photo micrograph illustrating precision 7.5 IR induced total salivary tissue volume reduction at necropsy compared to sham treated control young female mouse. **C**) X-Y plot showing precision IR dose dependent changes in the body weight adjusted wet weight of total salivary tissue in mg (male n=5 except n=4 for 7.5Gy IR dose; female n=5 or old female n=4 for each experimental group. **D**) Photomicrograph of stained parotid, submandibular and sublingual glands of young female NSG-GM3 mice at 6 month post-IR (0 and 7.5 Gy). Five micron sections were stained with (i) H&E, (ii) Sirius red, (iii) Trichrome, (iv) EpCAM (v) Aquaporin 5/beta-catenin and (vi) NKCC1 (female n=3). ns denotes not significant; **p<0.001.

The advantages of developing precision-IR based female NSG-GSM3 as a chronic hyposalivation model for preclinical testing are manifold. *Animal:* These mice express human interleukin-3 (hIL3), human granulocyte monocyte colony stimulating factor (hGM-CSF) and human c-Kit ligand also known as human stem cell factor (hSCF), which makes them ideal to study the regenerative properties of c-KIT-expressing putative human salivary stem cells (10). Male NSG-SGM3 mice, like other rodents (23, 29), have granular convoluted tubular structures which are absent in human. Both young and old female mice are suitable models, but young female mice will perhaps offer quicker test results. The earliest post-IR timepoint which indicates stabilization of flow rate fluctuations in these female mice are around 2 months post-irradiation. We recommend this as the earliest timepoint to assess chronic hyposalivation levels. *Precision IR:* The presence of *Prkdc^scid^* mutation makes these mice radiosensitive due to the lack of functional DNA repair machinery (11). Head & neck radiation to overcome this issue may lead to cognitive and other brain defects (30). Therefore, precision IR enables hyposalivation studies using 5-7.5 Gy doses over chronic timeline irrespective of gender or age. These mice are available at Jackson Laboratories and breed well under standard conditions. *Schirmer’s test:* Each strip costs around 0.3US Dollar. Single kinetic measurement of a flow rate over 30 minutes in a single animal requires 6 strips. The reading can be noted real-time or a pencil mark is sufficient to analyze the strips later with no concerns of evaporation.

On the basis of our characterization of immunodeficient NSG-SGM3 mice following brain-sparing precision IR of salivary glands, we propose a novel chronic hyposalivation model for regenerative medicine and human cell therapy studies.

## Methods

### Animals

Female and male NOD.Cg-Prkdcscid Il2rgtm1Wjl Tg(CMV-IL3,CSF2,KITLG)1Eav/MloySzJ mice (8-12 weeks old) were purchased from Jackson Laboratories and maintained at Mayo Clinic animal facility. Experimental mice were housed in barrier facility and fed ad libitum with food pellets and acidified water. All procedures were approved by Mayo Clinic’s Institutional Animal Care and Use Committee (IACUC).

### Salivary targeted precision IR set-up

Each mouse was anesthetized via isoflurane for the duration of the procedure via a nose cone delivery system on a bite block, and placed in a Feet First Prone (FFP) position in the irradiation chamber. A commercially-available stereotactic stage (Model 900M, Kopf Instruments, Tujunga, CA) was modified to facilitate mounting to the X-RAD SmART irradiator (31) (Fig1A). X-ray irradiation (0, 2.5, 5 or 7.5 Gy; at ~5Gy/min dose rate) using X-RAD SmART irradiator was delivered to head and neck in a brain-sparing manner. The stereotactic X-RAD SmART irradiator set-up doesn’t require additional blocking. In brief, each mouse was volumetrically imaged with CBCT (60 kVp, 0.3 mA, 2.0 mm Al filter, 256 projections 0.2 mm^3^ voxel size). Subsequently, the filter was switched to high energy copper (irradiation) filter and beam is focused on the target using 10 mm circular collimator for brain-sparing salivary-ablative radiation treatment. Each mouse was under isoflurane anesthesia for about 10 min to complete the irradiation procedure.

### Pilocarpine-stimulated reflex saliva flow measurements

Animals were anesthetized using 2% isoflurane and injected sub-cutaneously with 2 mg/kg pilocarpine, a cholinergic drug. After 5 minutes, animals were restrained and the secreted saliva was collected continuously from the floor of the mouth using pre-weighed Schirmer’s test strips (Tearflo, HUB Pharmaceuticals LLC, Rancho Cucamonga, CA, USA) for a period of 30 minutes. The strips were changed every five minutes and the distance of saliva migration under the capillary action (mm) was recorded. Subsequently, the weight of blotted strips was determined gravimetrically. Saliva was measured every three weeks interval in all groups of mice for a period of 24 weeks. The rate of saliva secretion by Schirmer’s test and gravimetry was determined in irradiated and sham treated animals.

### Histology and immunostaining

Mouse major salivary glands were fixed in 10% formalin buffer fixative (fisher scientific; SF-1004) for 1 h prior to processing. Paraffin-embedded salivary glands tissues sections (5μm thick) were stained with Hematoxylin and Eosin (H&E; morphological changes), trichrome, and red stain (collagen). Stained slides were photographed using color brightfield imaging with Cytation5 (BioTek Instruments Agilent Technologies, USA). For immunostaining, 5 μm sections salivary tissues were deparaffinized with xylene, rehydrated in graded ethanol and rinsed with distilled water. Slides were rehydrated and antigens retrieved by boiling slides for 20 minutes in antigen unmasking solution (Vector Laboratories Inc, Burlingame, CA, USA). Sectioned tissues were incubated overnight at 4°C with desired primary antibody anti-EpCAM antibody conjugated with Alexa Fluor 488, (Abcam, ab237384 1:200 dilution), anti-NKCC1(Abcam, ab59791; 1;250), anti-Aquaporin-5 (Sigma-Aldrich, AB15858, 1:200), or anti-β catenin conjugated with Alexa Fluor 647 (Abcam, ab194119; 1;250) diluted in blocking buffer (2% normal goat serum, 0.1% Triton X-100 in PBS). After three washes in PBS-Tween 20 (PBST), sections were incubated for 1 h with secondary antibody (Alexa Fluor® 488 goat anti-rabbit IgG (ThermoFisher Scientific, A-11008, 1:200), or Alexa Fluor^®^ 594 goat anti-rabbit IgG (ThermoFisher Scientific, A-11012, 1:200) diluted in PBS, containing 2% normal goat serum. Nuclei were counterstained with DAPI Fluoromount-G^®^ (Southern Biotech, Birmingham, AL, USA).

### Statistical Analysis

Analyses were performed using GraphPad Prism 8.0. For parameteric comparisons between treated and untreated groups, an unpaired two-tailed Student’s t-test was utilized. P-values of < 0.05 were considered significant. Multiple group comparison was performed by one-way or two-way ANOVA. Differences were considered significant at P < 0.05. Results are reported as mean ± SEM unless otherwise stated.

## Contributions

SA and IV conducted experiments and analyzed the results, performed statistical analysis, and wrote the manuscript. ET, MW and JS assisted with SmART technology and stereotactic radiotherapy design and dosimetry. JS, XX and CE assisted with data collection. RS and JG assisted with interpretation of histology and immunohistochemistry data. IV performed statistical analysis. IL provided insight on current saliva collection methods. JJ and NK conceptualized and designed the study, interpreted the data, and wrote the manuscript. NK supervised the study. All authors contributed to the drafting of the manuscript.

## Acknowledgments

This study was supported by Mayo Clinic and in part by grants to JJ and NK from Mayo Clinic Center for Regenerative Medicine. We thank Dr. James E. Melvin (NIH/NIDCR) for helpful discussions on salivary measurement methods, and Drs. Geng Xian Shi and Gang Liu for their technical assistance.

