## Supplementary Information for "Distinct Long-term Effects of Precision X-Radiation on Reflex Saliva Flow Rate and Tissue Integrity in a Preclinical Model of Chronic Hyposalivation"

**Supplementary figures and tables**

**Supplementary figure 1: Histological evaluation of submandibular gland in sham irradiated male (A) and female (B) NSG-SGM3 mouse at postnatal day 270**. The arrowhead denotes granular convoluted tubule (GCT); scale bar- 200µm. The GCT occupies a relatively large proportion of the glandular tissue and the cells are packed with dark secretory granules in submandibular glands of male compared to female which contain less GCTs.

**Supplementary table 1:** Depicting the summary of methods reported for modelling radiation induced hyposalivation and measurement of salivary flow rates in various animal models.

Supplementary Figure-1

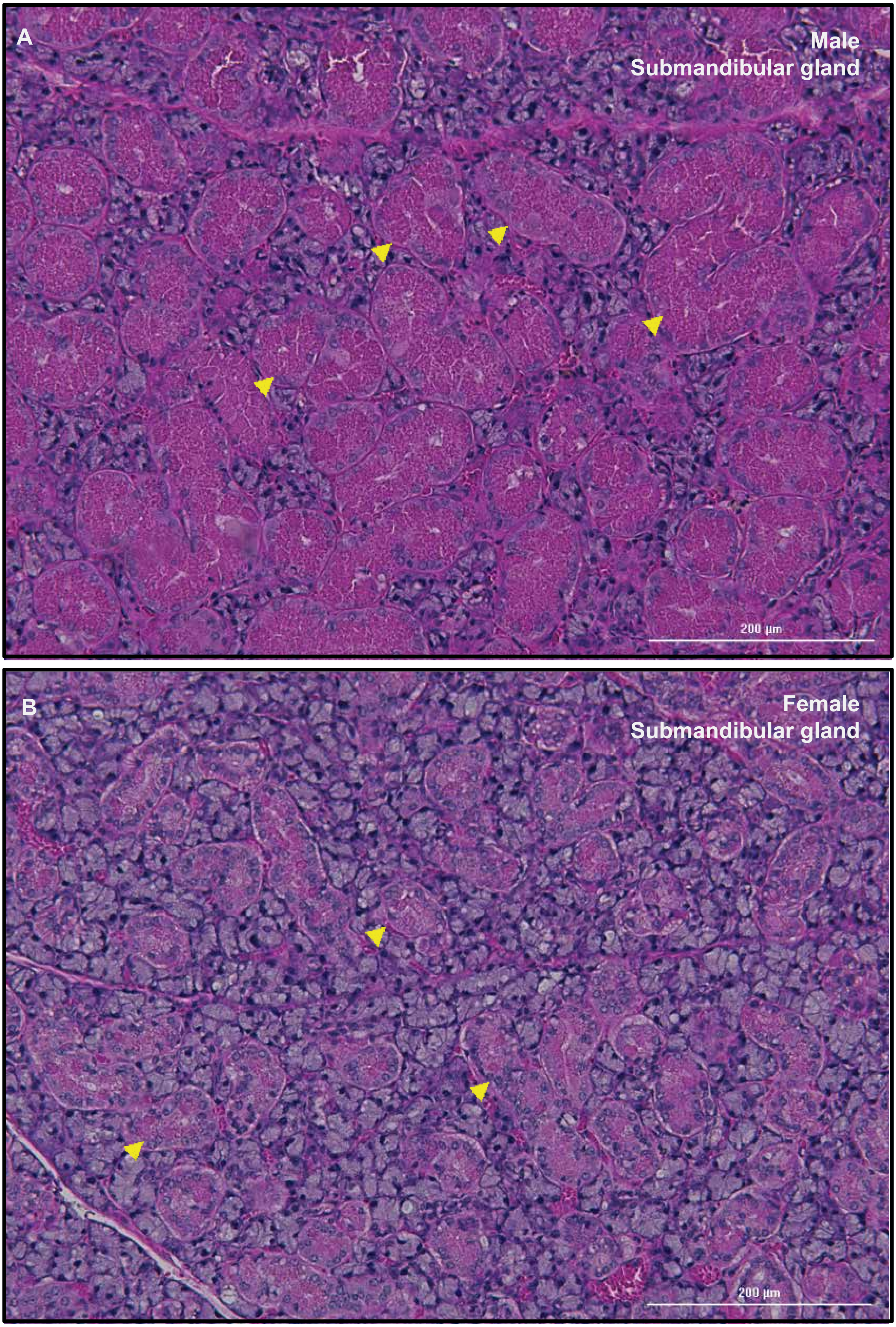

**Supplementary Table 1.** Summary of methods reported for modelling radiation induced hyposalivation in animals.

| Animal Model | Age (wk) | Gender | Dose (Gy) | Type of radiation | Anatomical site | Anesthesia | Stimulation | Method | Length (mins) | Immuno-deficient? | Xeno-transplantable? | Ref |
|---|---|---|---|---|---|---|---|---|---|---|---|---|
| **Mouse** | | | | | | | | | | | | |
| **C57BL/6** | 4-8 | Female | 5 (SD) | γ- rays | H&N | Ketamine/Xylazine | Carbachol | Gravimetry | 5 | No | No | (1) |
| | 8-10 | Female | 15 (SD) | γ- rays | SMG | Ketamine/Xylazine | Pilocarpine | Gravimetry | 10 | No | No | (2) |
| | 10-12 | Female | 15 (SD) | X- rays | SMG | Ketamine/Xylazine | Pilocarpine | Gravimetry | 15 | No | No | (3) |
| | | | 0-15 | X- rays | SG & oral mucosa | | Carbomylcholine chloride | Gravimetry | 4 | No | No | (4) |
| | 5 | Female | 25 | X- rays | SMG | | Pilocarpine | Gravimetry | 10 | No | No | (5) |
| | 8 | | 18 | X- rays | SMG | Sodium pentobarbital | Pilocarpine | Gravimetry | 15 | No | No | (6) |
| | 4-5 | Female | 15 (SD) 30(SpD) | | SMG& UN | | Pilocarpine | Gravimetry | | No | No | (7) |
| | 6 | Male | 15 (SD) | X- rays | SMG | Ketamine/Xylazine | Pilocarpine | Gravimetry | 15 | No | No | (8) |
| | 8-12 | Female | 15 (SD) | X- rays | SG | | Pilocarpine | Gravimetry | 15 | No | No | (9) |
| | 9-12 | Female | 15 (SD) | X- rays | SG | | Pilocarpine | Gravimetry | 15 | No | No | (10) |
| | 4-5 | Female | 15 (SD) | | SG | | Pilocarpine | Gravimetry | | No | No | (11) |
| | | | 15 (SD) | X- rays | H&N | | Pilocarpine | Gravimetry | | No | No | (12) |
| **C3H** | 7-9 | Female | 15 (SD) | X- rays | H&N | Ketamine/Xylazine | Pilocarpine | Gravimetry | 10 | No | No | (13) |
| | 7-9 | Female | 15 (SD) | | H&N | Ketamine/Xylazine | Pilocarpine | Gravimetry | 10 | No | No | (14) |
| **FVB/NJ** | 4-8 | Female | 5 (SD) | γ- rays | H&N | Ketamine/Xylazine | Carbachol | Gravimetry | 5 | No | No | (1) |
| **FVB** | 4-5 | Female | 5 (SD) | γ- rays | SG | | Carbachol | Gravimetry | 5 | No | No | (15) |
| **ICR** | 5 | Female | 25 | X- rays | SMG | | Pilocarpine | Gravimetry | 10 | No | No | (5) |
| **ICR-nu/nu** | 5 | Female | 25 | X- rays | SMG | | Pilocarpine | Gravimetry | 10 | Yes | No | (5) |
| **SCID** | 4-5 | Female | 15 (SD) 30(SpD) | | SMG& UN | | Pilocarpine | Gravimetry | | Yes | Yes | (7) |
| **NSG** | | | 5 (SD) | | SG | | Pilocarpine | Gravimetry | 15 | Yes | Yes | (16) |
| **Rats** | | | | | | | | | | | | |
| **Sprague-Dawley** | | Male | 15 (SpD) 75 (TD) | | Left mandible | Ketamine/Medetomidine | Pilocarpine | Gravimetry | 15 | No | No | (17) |
| | | | 15 | | SMG/SLG | | Pilocarpine/ Cevimeline | | | | | (18) |
| | 7-8 | Male | 15 | X- rays | H&N | | Pilocarpine | Gravimetry | 120 | No | No | (19) |
| **Wistar** | | Male | 30 | γ- rays | Left parotid | Ketamine/Xylazine | Pilocarpine | Gravimetry | 40 | No | No | (20) |
| | | Male | 20 | | SMG | | | Gravimetry | | No | No | (21) |
| | 8-9 | Male | 10,15,20 (SD) | X- rays | Right parotid | Isoflurane/$O_2$ | Pilocarpine | Gravimetry | 30 | No | No | (22) |
| | | | 15 (SD) | γ- rays | | | Norepinephrine | | | | | (23) |
| **Rabbits** | | | | | | | | | | | | |
| **Chinchilla Bastard** | | Female | | X- rays | Parotid SMG | | Carbachol | Sialometry - scintigraphy | 25 | No | No | (24) |
| **Pigs** | | | | | | | | | | | | |
| **Yucatan miniature** | ~10 | | 10 (SD) | | Right parotid | Ketamine/Xylazine Isoflurane/$O_2$ | Pilocarpine | Gravimetry | 10 | No | No | (25) |
| **BA-MA miniature** | 8 mnth | Male | 20 | | Right parotid | Ketamine/Xylazine | Pilocarpine | Gravimetry | 10 | No | No | (26) |

Wk- weeks, SD- single dose, SpD- Spilt dose, TD- total dose, H&N- head and neck, UN- upper neck, SG-salivary gland, SMG –submandibular gland, SLG-sublingual gland
